# VPS35/Retromer-dependent MT1-MMP regulation confers melanoma metastasis

**DOI:** 10.1101/2025.02.25.640017

**Authors:** Qinggang Hao, Yan Bai, Ruiqi Guan, Rui Dong, Weiyu Bai, Hayam Hamdy, Liqiong Wang, Mingyao Meng, Yan Sun, Junling Shen, Jianwei Sun

## Abstract

Retromer is a conserved endosomal trafficking complex responsible for recycling transmembrane protein cargoes. Membrane-type I matrix metalloproteinase (MT1-MMP), a well-studied membrane-type metalloprotease, is highly expressed in metastatic melanomas. Previously, we reported that inducing MT1-MMP perinuclear localization and inhibiting MT1-MMP membrane localization significantly reduce melanoma metastasis. However, the regulation of MT1-MMP subcellular localization and recycling is still largely unknown. Here, we performed target gene shRNA screening and found that shRNA targeting the retromer complex subunit vacuolar protein sorting 35 (VPS35) inhibited MT1-MMP membrane localization and induced its perinuclear localization. We found that inhibiting VPS35/retromer decreased MT1-MMP recycling and increased MT1-MMP-lysosome localization, which significantly affected the stability of MT1-MMP. Furthermore, our results indicated that VPS35/retromer regulates the transcription of *MT1-MMP* through the activation of the IL6/STAT3 inflammatory signaling pathway. Tissue microarray analysis indicated that VPS35/retromer positively correlated with MT1-MMP levels and distant metastasis. Xenograft experiments showed that targeting VPS35/retromer significantly inhibited melanoma lung metastasis, which is dependent on MT1-MMP. Our results implicate the importance of VPS35/retromer in metastatic dissemination. Our study suggests that targeting the VPS35/retromer-MT1-MMP axis will contribute to inhibiting the metastasis of melanoma.

## INTRODUCTION

Cancer metastasis refers to the spread of cancer cells from primary tumors to distant organs, accounting for 90% of cancer-related patient deaths (Chaffer and Weinberg, 2011). Despite significant advances in the treatment of primary tumors, managing tumor metastasis in advanced-stage cancer remains a major clinical challenge. The targeted degradation of the extracellular matrix (ECM) by invasive cells is crucial for cancer cell metastasis. Proteolysis of the ECM is a key factor in tumor cell metastasis.

ECM proteolysis is critical for cancer cell invasion and metastasis during cancer progression. Matrix metalloproteinases (MMPs) are major contributors to the proteolytic degradation of the ECM during tumor invasion. Membrane-type I matrix metalloproteinase (MT1-MMP/MMP14) is a well-characterized membrane-bound metalloprotease and akey driver of ECM degradation. *MT1-MMP* is highly expressed in metastatic cancers. It is associated with poor prognosis, and has emerged as a proven clinical target. Several research groups have developed selective MT1-MMP inhibitory antibodies targeting either the catalytic domain or the collagen-binding domains, which have been shown to effectively limit tumor growth and metastasis in triple-negative breast cancer (TNBC) tumor growth and metastasis in xenograft assays(Ager et al., 2015; Botkjaer et al., 2016; Devy et al., 2009). MT1-MMP undergoes continuous trafficking and recycling within the cell. It moves between different subcellular regions and circulating on the cell surface, thereby maintaining its activity in ECM degradation and tumor invasion. This process is a crucial regulatory factor in tumor invasion and metastasis(Loskutov et al., 2015).

The retromer complex is an evolutionarily conserved protein complex that regulates endosomal protein sorting and recycling, and it plays a crucial role in intracellular transport processes such as developmental signal transmission, polarized transport, and lysosome biogenesis (Haft et al., 2000; Harterink et al., 2011; Seaman, 2021). Vacuolar protein sorting 35 (VPS35), a vital component of the retromer, significantly influences several cancer types, including breast cancer, liver cancer, and melanoma(Trousdale and Kim, 2015). It regulates tumor occurrence by modulating the N-Ras signaling pathway, facilitating the retrograde transport of N-Ras from endosomes to the Golgi and plasma membrane (Goodwin et al., 2005; Rocks et al., 2005). *VPS35* also impacts the Wnt signaling pathway. It is critical for cell migration, as its dysregulation results in Wnt/PCP pathway defects driving tumor metastasis (Butler and Wallingford, 2017; Caddy et al., 2010; VanderVorst et al., 2018). VPS35/retromer-dependent endosomal recycling of MT1-MMP is linked to cancer metastasis and poor prognosis. Targeting the retromer inhibits MT1-MMP recycling to the cell membrane. Recently, it has been reported that the retromer regulates MT1-MMP recycling in breast cancer (Sharma et al., 2020). However, its role in melanoma and the regulation of *MT1-MMP* by the retromer remain largely unknown.

Epidemiological studies highlight chronic infections and inflammation as significant risk factors for various cancer types. *STAT3* is a key oncogenic gene that promotes tumor development and it is essential for cell transformation mediated by oncogene such as Src, which are directly affected in human cancers(Bowman et al., 2001). Additionally, *STAT3* suppresses anti-tumor immunity by antagonizing the expression of T helper 1 cytokines, including IL-12 and interferon-γ, which are crucial for both innate and T cell-mediated anti-tumor responses(Rébé and Ghiringhelli, 2019).

*STAT3* often synergizes with other signaling pathways, such as NF-κB, a critical regulator of inflammation and carcinogenesis. It interacts with *STAT3* to control the expression of genes involved in tumor progression(Zou et al., 2020). IL-6 promotes the invasion and metastasis of tumor cells by activating STAT3. Furthermore, *STAT3* promotes ECM degradation by regulating the expression of MMPs, such as MMP-2 and MMP-9, thus facilitating tumor cell invasion into adjacent tissues and blood vessels, and driving metastasis(Hu et al., 2024).

In this study, we identified VPS35 a key subunit of a retromer complex, as a critical regulator of MT1-MMP membrane localization and VPS35 also induced its perinuclear localization. Silencing of VPS35/retromer decreased MT1-MMP recycling and increased its lysosomal localization, thus significantly affecting MT1-MMP protein stability. Additionally, our results further demonstrated that VPS35/retromer regulates *MT1-MMP* mRNA levels via the activation of the IL6/STAT3 inflammatory signaling pathway. Tissue microarray analysis revealed a positive correlation between VPS35/retromer expression and *MT1-MMP* levels, and distant metastasis. In vivo, xenograft experiments confirmed that retromer inhibition significantly reduced melanoma lung metastasis in an MT1-MMP-dependent manner. Collectively, these findings underscore the critical role of the retromer-MT1-MMP axis in melanoma metastasis and suggest that targeting this pathway could represent a promising therapeutic strategy for inhibiting metastatic progression.

## RESULTS

### VPS35/Retromer is involved in MT1-MMP trafficking and stability

In our previous studies, we found that targeting MT1-MMP recycling or inducing MT1-MMP perinuclear localization significantly inhibits melanoma metastasis(Sun et al., 2014). However, the regulation of MT1-MMP subcellular localization and recycling remains largely unknown. To explore the mechanism of MT1-MMP recycling and its role in cancer metastasis, we performed target gene screening focusing on protein transport.

We first constructed a stable melanoma WM793 cell line expressing MT1-MMP-GFP. We screened for genes involved in intracellular vesicle transport. We examined the effects of Rabs, VAMP7, and retromer components on the subcellular localization of MT1-MMP. The result showed that knocking down *VPS35*, a retromer component, significantly reduced the membrane MT1-MMP-GFP signal, suggesting that VPS35/retromer may regulate MT1-MMP recycling in melanoma **(Figure 1A)**. *VPS35* shRNA also significantly increased the percentage of cells with MT1-MMP cytoplasmic aggregates **(Figure 1B and C)**. To demonstrate that *VPS35* shRNA decreases membrane localization of MT1-MMP by regulating its recycling, we performed a biotinylation assay and found that *VPS35* shRNA significantly reduced membrane-localized MT1-MMP **(Figure 1D)**. Interestingly, *VPS35* shRNA not only affected MT1-MMP subcellular localization, but also significantly reduced the total MT1-MMP level **(Figure 1D and E)**.

**Figure 1.**
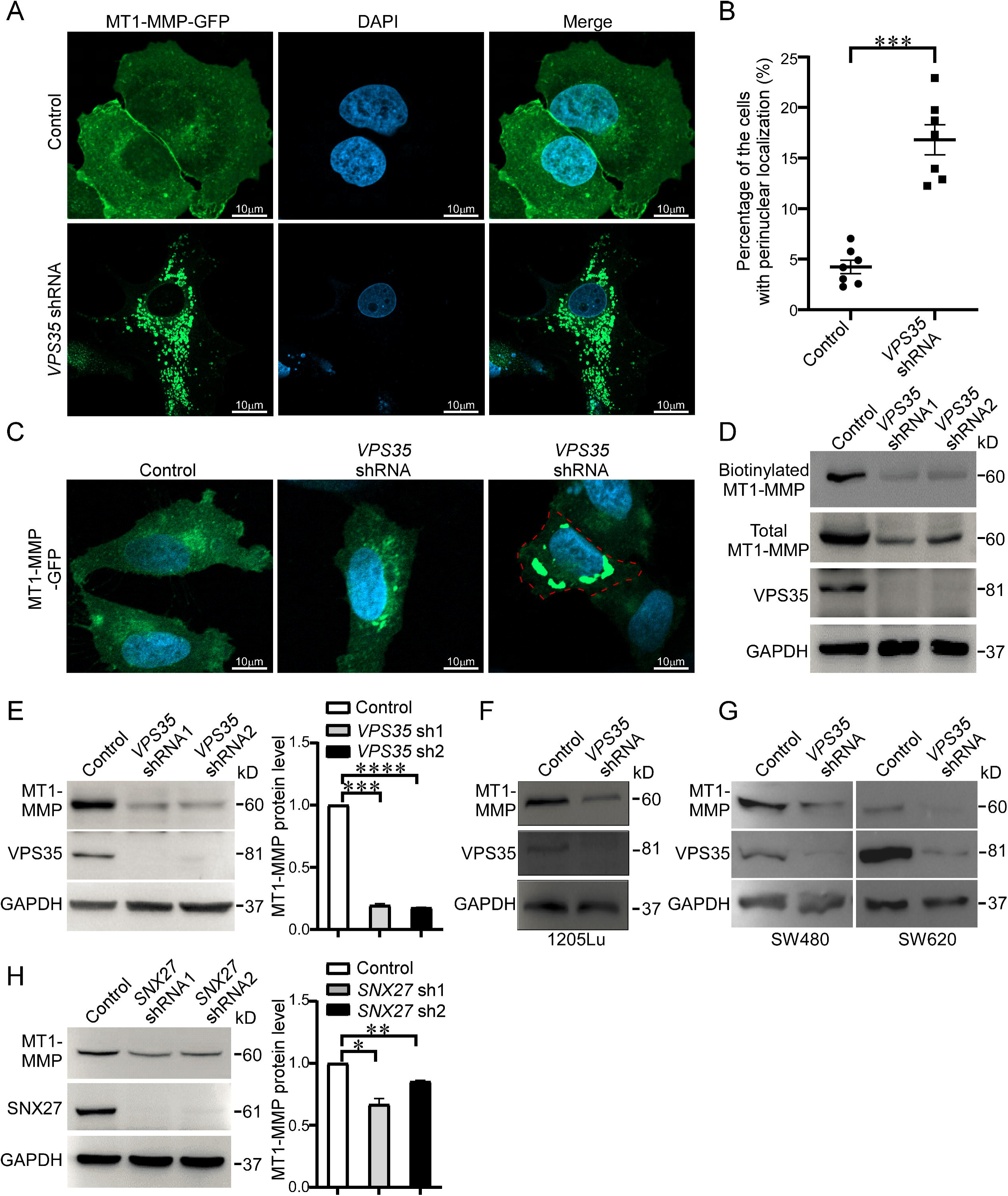
VPS35/Retromer is involved in MT1-MMP trafficking and stability. A. Fluorescent microscopy images showing the localization of MT1-MMP-GFP in control cells versus *VPS35*-knockdown cells. B. Percentage of cells with perinuclear localization of MT1-MMP-GFP in *VPS35* shRNA cell. C. Fluorescent microscopy images showing the cell membrane localization of MT1-MMP-GFP in control cells versus *VPS35*-knockdown cells. D. Biotin assay showing a decrease in membrane-localized MT1-MMP. E. *VPS35* shRNAs reduces total MT1-MMP levels in WM793 cells. F. *VPS35* shRNA reduces total MT1-MMP levels in 1205Lu cells. G. *VPS35* shRNA reduces total MT1-MMP levels in SW480 and SW620 colon cells. H. *SNX27* shRNA reduces total MT1-MMP levels in WM793 cells. (\**P* < 0.05, \*\**P* < 0.01, \*\*\**P* < 0.001, \*\*\*\**P* < 0.0001)

To further verify that VPS35 affects MT1-MMP levels, we knocked down *VPS35* in 1205Lu melanoma cells and two colon cancer cell lines, SW480 and SW620. We found that *VPS35* shRNA significantly reduced MT1-MMP levels in these cells **(Figure 1F and G)**. Additionally, we knocked down *SNX27*, another component of the retromer complex, and observed a similar reduction in MT1-MMP levels **(Figure 1H)**. This suggests that *VPS35* shRNA not only reduces membrane-localized MT1-MMP, but also decreases the total MT1-MMP level.

*VPS35* shRNA affected the membrane localization of MT1-MMP, suggesting that VPS35 may play a role in tumor metastasis. To further determine the role of VPS35 in tumor progression, we analyzed its expression levels in 33 cancer types using the The Cancer Genome Atlas (TCGA) and Genotype Tissue Expression Project (GTEx) databases. *VPS35* is highly expressed in almost all tumors, particularly in colorectal and pancreatic cancers, compared to normal tissues (**Figure S1A**). In breast invasive carcinoma, brain lower grade glioma, lung squamous cell carcinoma, and stomach adenocarcinoma, high VPS35 levels were significantly negatively correlated with patient survival rates (**Figure S1B-E**), suggesting that VPS35 is associated with cancer progression.

### VPS35/Retromer affects MT1-MMP protein stability through the lysosome

To investigate the mechanism by which VPS35/retromer regulates MT1-MMP levels, we first assessed MT1-MMP stability in control and *VPS35* shRNA. Using the 10 μg/mL protein synthesis inhibitor cycloheximide (CHX) to treat both control and *VPS35* shRNA cells, we found that MT1-MMP degraded more rapidly in *VPS35* shRNA cells compared to control cells (**Figure 2A-D**). Intracellular protein degradation can occur via the ATP-dependent proteasome pathway or the ATP-independent lysosomal pathway. To examine the degradation mechanism of MT1-MMP by *VPS35* shRNA, we treated the cells with the 5 μM proteasome inhibitor MG132 and the 100 nM lysosomal inhibitor Bafilomycin A1 (BFA1, inhibiting lysosomal degradation) for 6 hours, respectively. Notably, BFA1 treatment increased MT1-MMP levels in both WM793 and *VPS35* shRNA cells compared to the untreated and MG-132 treated groups, indicating that BFA1 exerts an inhibitory effect on MT1-MMP degradation in *VPS35* shRNA cells (**Figure S2A, Figure 2E and F**). In contrast, MG-132 had a minimal effect on MT1-MMP levels, suggesting that the proteasomal pathway plays a minor role in this context. These findings suggest that the rapid degradation of MT1-MMP caused by *VPS3*5 shRNA occurs primarily via the lysosomal pathway.

**Figure 2.**
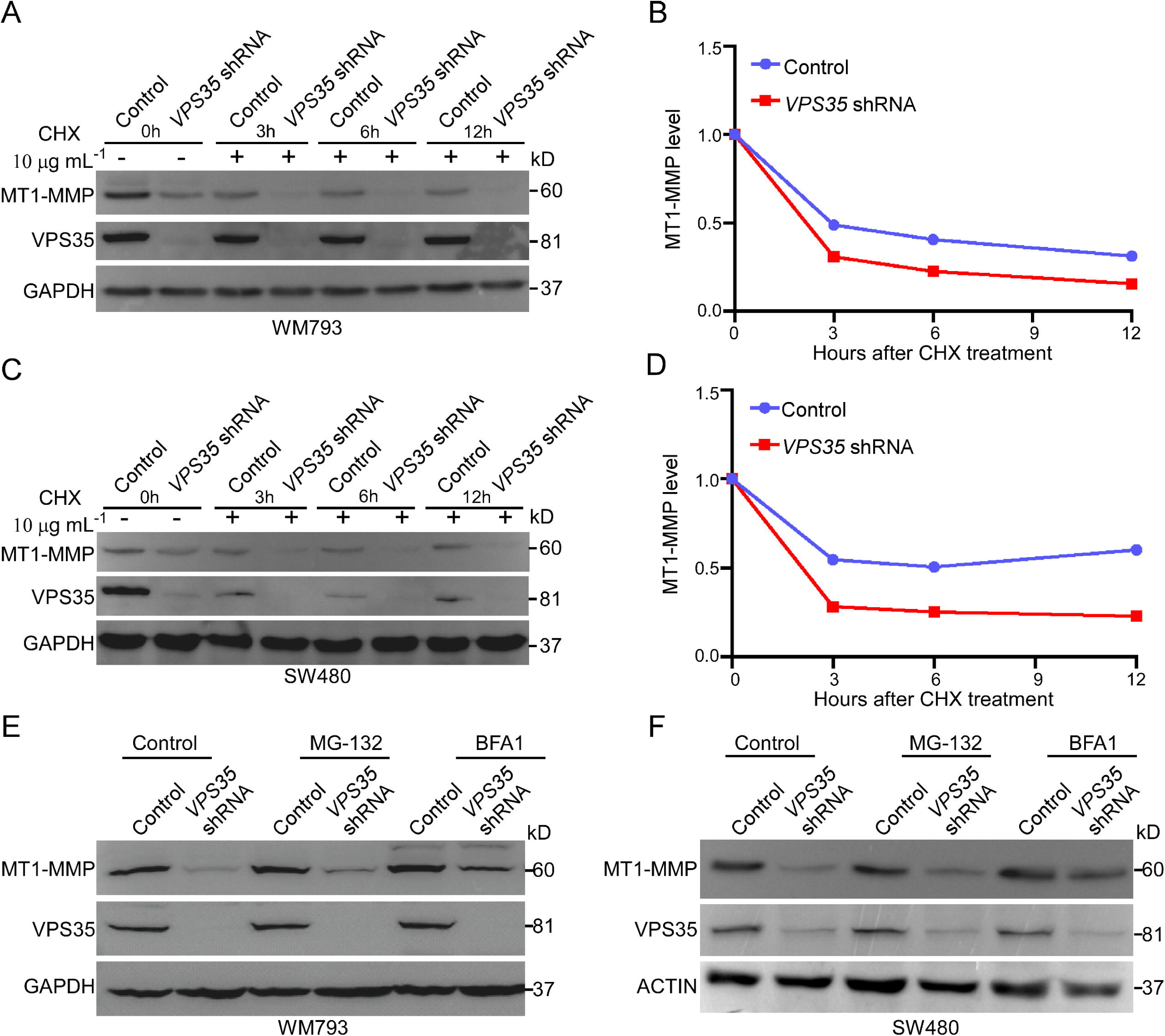
VPS35/Retromer affects MT1-MMP protein stability through the lysosome pathway. A. Western blotting analysis of MT1-MMP stability following 10 μg/mL CHX treatment in WM793 control and *VPS35* shRNA cells. B. Quantitative analysis of MT1-MMP stability in WM793 control and *VPS35* shRNA cells. C. Western blotting analysis of MT1-MMP stability following 10 μg/mL CHX treatment in control and *VPS35* shRNA SW480 cells. D. Quantitative analysis of MT1-MMP stability in control and *VPS35* shRNA SW480 cells. E. Western blotting analysis of MT1-MMP levels following 5 μM MG-132 and 100 nM BFA1 treatment in WM793 control and *VPS35* shRNA cells. F. Western blotting analysis of MT1-MMP levels following 5 μM MG-132 and 100 nM BFA1 treatment in SW480 control and *VPS35* shRNA cells.

### Inhibition VPS35 increased the lysosomal localization of MT1-MMP

To explore the mechanism by which VPS35 regulates MT1-MMP recycling and stability, we first confirmed whether VPS35 and MT1-MMP form a complex. Co-IP results suggest that MT1-MMP was an interacting partner of VPS35 (**Figure 3A**). The immunofluorescence results showed shat the colocalization between VPS35 and MT1-MMP and found strong colocalization in the cells (**Figure 3B**). Proximity Ligation Assay (PLA) is a highly sensitive technique that quantitatively analyzes protein-protein interactions within cells by detecting the spatial proximity between proteins. As shown in Figure 3C, the binding of MT1-MMP and VPS35 was located in the cytoplasm of tumor cells (**Figure 3C**). These results suggest that MT1-MMP could be recycled by the VPS35/retromer complex.

**Figure 3.**
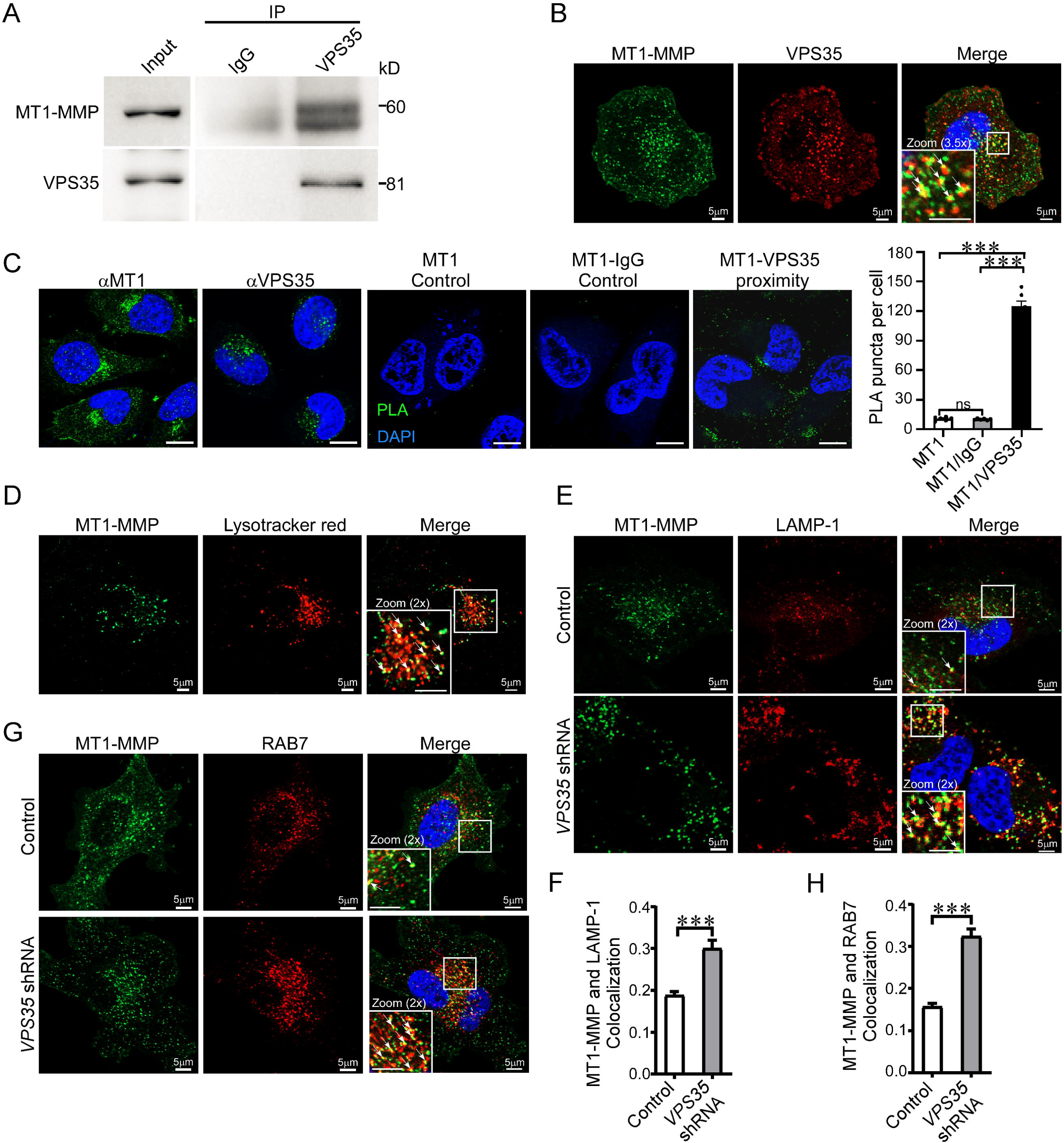
Inhibition of VPS35/retromer increases MT1-MMP colocalization with lysosomes. A. Co-IP analysis shows that VPS35 interacts with MT1-MMP. B. Co-localization analysis of MT1-MMP and VPS35, scale bar, 5 μm. C. PLA analysis co-localization between VPS35 and MT1-MMP, scale bar, 10 μm. D. Co-localization analysis of MT1-MMP and the lysotracker, scale bar, 5 μm. E. Co-localization analysis of MT1-MMP and the lysosomal marker LAMP-1 in control and *VPS35* shRNA cells, scale bar, 5 μm. F. Statistical analysis of the co-localization rate between MT1-MMP and the lysosomal marker LAMP-1 in control and *VPS35* shRNA cells. G. Co-localization analysis of MT1-MMP and the late endosome marker RAB7 in control and *VPS35* shRNA cells, scale bar, 5 μm. H. Statistical analyzation of the co-localization rate between MT1-MMP and RAB7 in control and *VPS35* shRNA cells. (arrows indicate colocalization, \*\*\**P* < 0.001)

MT1-MMP degradation caused by *VPS35* shRNA occurs primarily via the lysosomal pathway. To test our hypothesis, we analyzed the co-localization of MT1-MMP and lysosomes in control and *VPS35* shRNA cells. Immunofluorescence showed that MT1-MMP co-localizes with lysosomes (**Figure 3D**). LAMP-1 is a marker for lysosomes, while RAB7 is a marker for late endosomes. The co-localization rate between MT1-MMP and the lysosome marker LAMP-1 increased in *VPS35* shRNA cells (**Figure 3E and F**). Retromer-mediated transmembrane protein recycling is intrinsically coupled to early-to-late endosome conversion along the endosome-lysosome fusion pathway. We found that *VPS35* shRNA also increases the co-localization rate between MT1-MMP and the late endosome marker RAB7 (**Figure 3G and H**). In sum, our results show that inhibition of VPS35/retromer increases the co-localization ratio between MT1-MMP and late endosomes and lysosomes, enhancing its degradation through lysosomes.

### VPS35/Retromer regulates MT1-MMP transcription through IL-6/STAT3

We further investigated whether VPS35/retromer affects MT1-MMP protein levels by influencing *MT1-MMP* transcription. Surprisingly, we found that *VPS35* shRNA significantly reduced *MT1-MMP* mRNA levels by approximately 40%, which was substantially less than the approximately 81% reduction observed at the protein level (**Figure 1E**, **Figure 4A**). Additionally, *SNX27* shRNA also reduced *MT1-MMP* mRNA levels (**Figure 4A**). To verify our results, we knocked down *VPS35* in 1205Lu cells, and the results showed that *VPS35* shRNA significantly reduced *MT1-MMP* mRNA levels (**Figure 4B**). These findings suggest that *VPS35* shRNA may affect the transcription of *MT1-MMP*.

**Figure 4.**
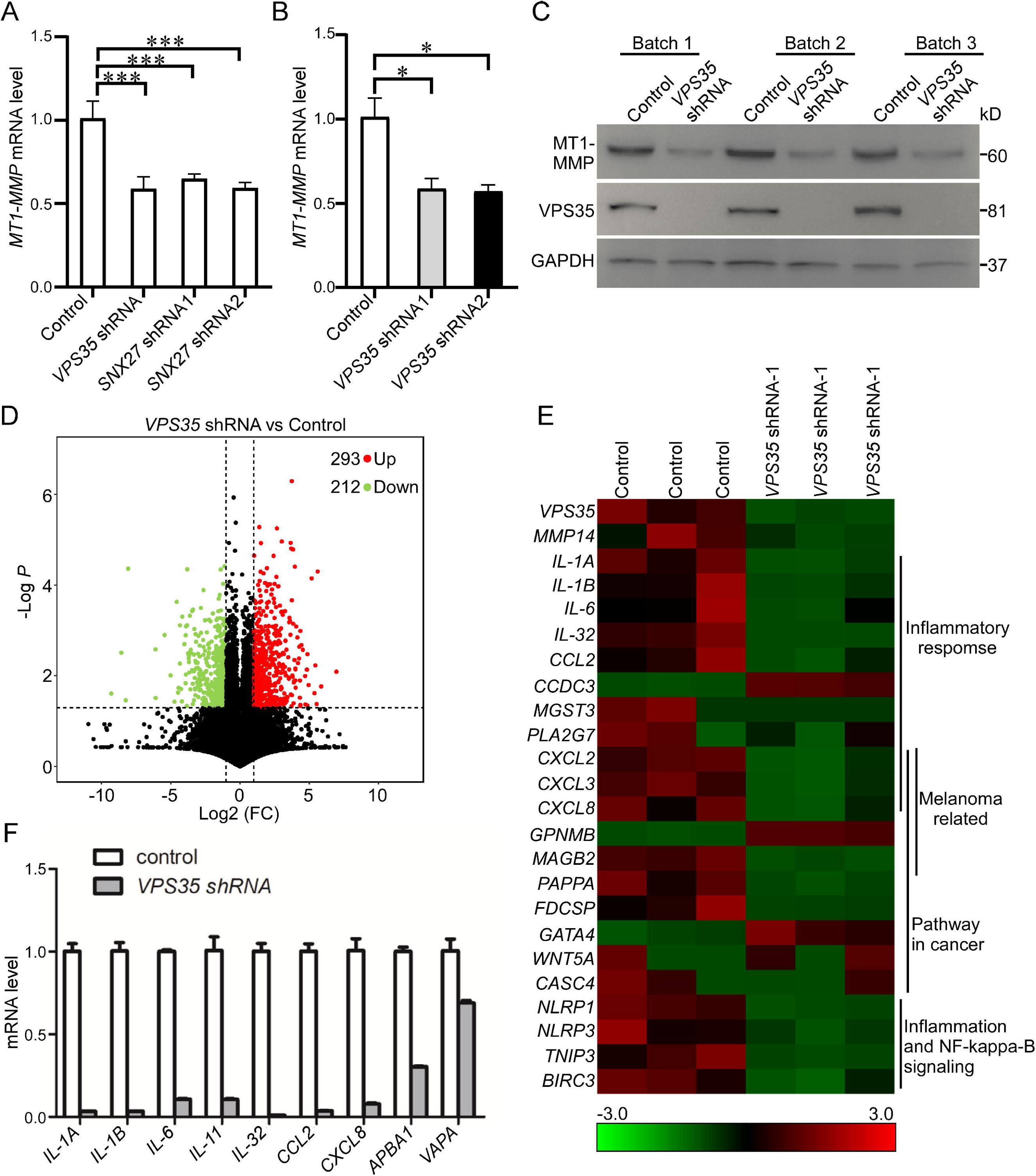
VPS35/Retromer regulates gene expression and inflammatory response. A. Effects of *VPS35* and *SNX27* shRNA on *MT1-MMP* mRNA levels in WM793 cells. B. Effects of *VPS35* shRNA on *MT1-MMP* mRNA levels in 1205Lu cells. C. Western blotting analysis of *VPS35* shRNA in three batches of cells. D. Volcano of differentially expressed genes in WM793 control and *VPS35* shRNA cells. E. Heatmap analysis of gene expression in WM793 control and *VPS35* shRNA cells. F. q-PCR analysis of differentially expressed genes in WM793 control and *VPS35* shRNA cells. (\**P* < 0.05, \*\**P* < 0.01, \*\*\**P* < 0.001)

To investigate how VPS35/retromer regulates *MT1-MMP* transcription, we performed RNA sequencing using WM793 control and *VPS35* shRNA cells (**Figure 4C**). Although the VPS35/retromer complex is not a transcription factor, our result showed that *VPS35* shRNA affects the mRNA levels of many genes (**Figure 4D**). Inflammatory factors play an important role in tumor metastasis, and our sequencing data revealed a significant reduction in the expression of numerous inflammatory factors. To verify the reliability of the sequencing results, we selected nine inflammatory genes for q-PCR validation in control and *VPS35* shRNA WM793 cells. The results showed that these nine genes were significantly reduced in *VPS35* shRNA cells, consistent with the sequencing data (**Figure 4F**). This indicates the sequencing quality is reliable. Further analysis revealed that inhibiting VPS35 significantly reduces the inflammatory response, melanoma-related genes, and genes involved in cancer pathways (**Figure 4E**). These findings suggest that VPS35 may regulate the transcription levels of downstream genes by modulating the inflammatory response.

Nuclear factor-kappaB (NF-κB) plays a crucial role in the development and progression of cancer. Inhibition of NF-κB activity significantly reduces the proliferation and invasion of Hep3B cells and down-regulates the expression of invasion-related molecules, including matrix metalloproteinase (MMP)-2, MMP-9, and MT1-MMP (Wu et al., 2009). Our KEGG and gene enrichment analyses show that inhibition of *VPS35* is closely related to the inflammatory response (**Figure 5A**). IL6/STAT3 inflammatory pathway plays a crucial role in tumor metastasis. To explore the mechanism by which VPS35 regulates inflammatory response, we investigated whether VPS35 and STAT3 form a complex. Co-IP analysis results showed that STAT3 co-precipitated with VPS35 (**Figure S2B**). The functions of *STAT3* and its intracellular activities are highly dependent on its phosphorylation status, specifically the activation level of p-STAT3 (Tyr705). Aberrant and persistent activation of STAT3 has been found in various types of cancers. When *VPS35* is inhibited, the level of p-STAT3 (Tyr705) significantly decreases (**Figure 5B**). IL-6 is the most important STAT3 activator in various tumors. We stimulated cells with IL-6-conditioned medium for 1.5 hours and observed a significant increase in p-STAT3 levels in the control group, confirming that IL-6 effectively activates the STAT3 pathway. In contrast, p-STAT3 levels were markedly reduced in the *VPS35* knockdown groups, indicating that *VPS35* knockdown inhibits STAT3 phosphorylation (**Figure 5C**). Additionally, we confirmed that *VPS35* shRNA significantly reduced nuclear localization of STAT3 (**Figure 5D**). In gastric cancer, inhibiting *VPS35* reduces IL-6 expression and STAT3 phosphorylation (Zhou et al., 2024). The retromer complex acts as a platform for STAT3 phosphorylation and can promote the nuclear localization of STAT3(Smalley et al., 2021). These results suggest that VPS35 mediates IL-6-induced STAT3 activation.

**Figure 5.**
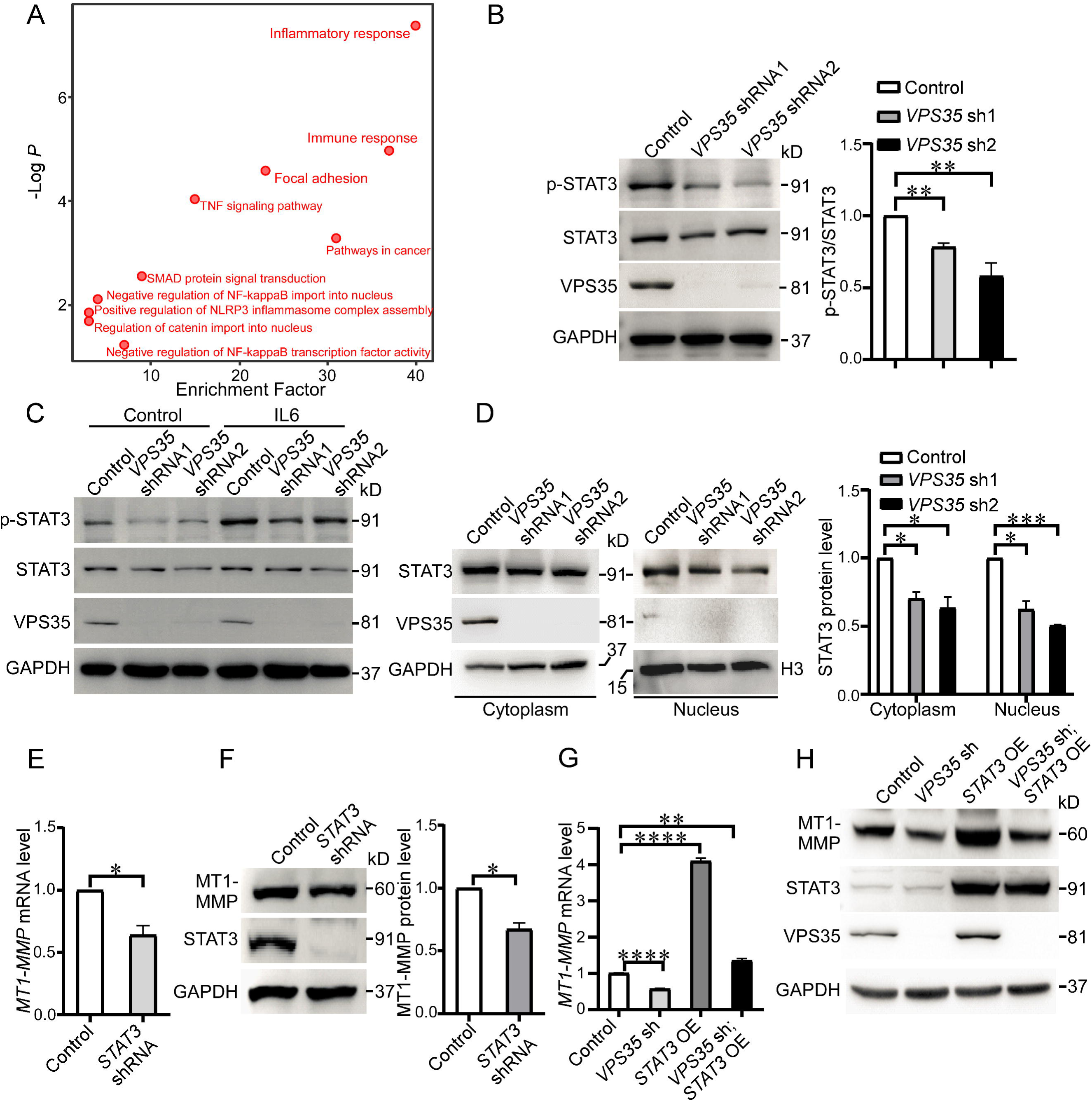
VPS35/Retromer regulates *MT1-MMP* transcription through IL-6/STAT3. A. KEGG enrichment analysis of differentially expressed genes in *VPS35* shRNA cells. B. Western blotting analysis of p-STAT3 (Tyr 705) levels in WM793 control and *VPS35* shRNA cells. C. Western blotting analysis of p-STAT3 (Tyr 705) levels after treating with IL6 in WM793 control and *VPS35* shRNA cells. D. Western blotting analysis and quantification of STAT3 nuclear localization in WM793 control and *VPS35* shRNA cells. E. q-PCR analysis of *MT1-MMP* mRNA levels in WM793 control and *STAT3* shRNA cells. F. Western blotting analysis of MT1-MMP protein levels in WM793 control and *STAT3* shRNA cells. G. q-PCR analysis of *MT1-MMP* mRNA levels following ectopic *STAT3* expression in *VPS35* shRNA cells. H. Western blotting analysis of MT1-MMP protein levels following ectopic *STAT3* expression in *VPS35* shRNA cells. (\**P* < 0.05, \*\**P* < 0.01, \*\*\**P* < 0.001, \*\*\*\**P* < 0.0001)

The IL-6/STAT3 signaling pathway constitutes a key oncogenic pathway and can drive cancer cell metastasis. *STAT3* and *MT1-MMP* are significantly co-expressed in melanoma (**Figure S2C**). To examine whether VPS35/retromer affects *MT1-MMP* transcription and tumor metastasis through the inflammatory IL-6/STAT3 signaling response, we knocked down *STAT3* in WM793 cells. *STAT3* shRNA significantly reduced *MT1-MMP* mRNA and protein levels (**Figure 5E and F**). We excluded the possibility that STAT3 regulates MT1-MMP protein levels through post-translational mechanisms. CHX treatment of *STAT3* shRNA cells revealed a gradual decrease in MT1-MMP levels over time in both control and *STAT3* knockdown groups, with no significant difference in degradation rates (**Figure S2D and S2E**). This indicates that *STAT3* knockdown does not accelerate MT1-MMP degradation. Furthermore, VPS35 levels remained unchanged in *STAT3* shRNA cells, suggesting *STAT3* does not regulate MT1-MMP via VPS35 (**Figure S2F**). These findings demonstrate that *STAT3* primarily regulates MT1-MMP at the transcriptional level. To verify whether VPS35 regulates *MT1-MMP* mRNA expression through *STAT3*, we ectopically expressed *STAT3* in *VPS35* shRNA WM793 cells. The results showed that ectopic expression of *STAT3* restored *MT1-MMP* levels in *VPS35* shRNA cells (**Figure 5G and H**). In conclusion, our results indicate that VPS35/retromer regulates *MT1-MMP* mRNA expression through the STAT3 inflammatory response.

### VPS35/Retromer regulates melanoma metastasis through *MT1-MMP*

To investigate if VPS35 regulates melanoma metastasis through MT1-MMP, we ectopically expressed *MT1-MMP* in *VPS35* shRNA 1205Lu melanoma cells (**Figure 6A and B**). Luciferase-labeled cells were injected via the tail vein into nude mice, and lung metastasis of 1205Lu melanoma cells was monitored using bioluminescence imaging (**Figure 6A**). Bioluminescence imaging and quantification of bioluminescence showed that knockdown of *VPS35* inhibited lung metastasis of melanoma and prolonged survival time of mice, but overexpression of *MT1-MMP* significantly restored tumor metastasis in *VPS35* shRNA 1205Lu melanoma cells and significantly reduced the mouse survival (**Figure 6C and D**). The inhibition of melanoma lung metastasis by *VPS35* shRNA was further confirmed by hematoxylin and eosin staining of the lungs harvested from nude mice **(Figure 6E)**. Collectively, our data suggest that VPS35/retromer-mediated regulation of MT1-MMP is critical for melanoma metastasis.

**Figure 6.**
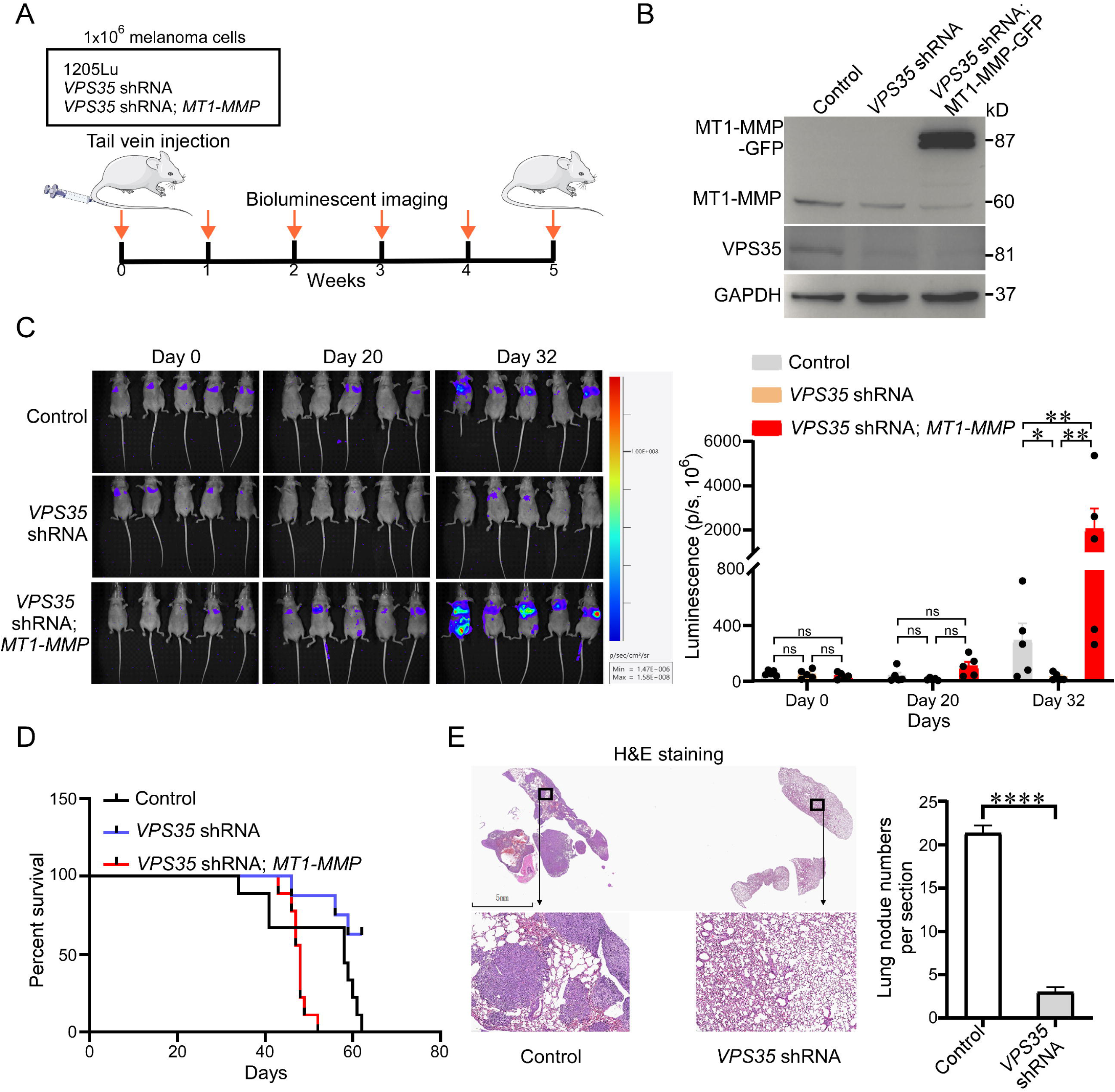
VPS35/Retromer promotes melanoma metastasis through MT1-MMP. A. Melanoma lung metastasis model via mouse tail vein injection. B. Western blotting analysis VPS35 and MT1-MMP in 1205Lu cells. C. *In vivo* bioluminescence imaging and quantification of bioluminescence (luminescence intensity in p/s, ×10^6^) shows pulmonary metastasis of melanoma in mice. D. Overall survival curves of melanoma lung metastasis mouse model. E. Hematoxylin and eosin (H&E) staining showed that *VPS35* shRNA in 1205Lu melanoma cells reduced metastasis in nude mice. (\**P* < 0.05, \*\**P* < 0.01, \*\*\*\**P* < 0.0001)

To evaluate the clinical significance of VPS35 in melanoma progression, we examined VPS35 and MT1-MMP levels in a melanoma tissue microarray. We found that VPS35 expression was significantly higher in patients with Clark IV and V melanoma compared to those with Clark II and III (**Figure 7A and B**). Prognostic analysis revealed that lower VPS35 levels were associated with longer overall survival (**Figure 7C**). Consistent with VPS35 expression, MT1-MMP levels also increased significantly with the progression of melanoma from Clark II to Clark V stage (**Figure 7D**). Additionally, we analyzed the correlation between VPS35 and MT1-MMP levels in the melanoma tissue microarray and the TCGA database. The data showed a significant positive correlation between *VPS35* and *MT1-MMP* mRNA levels in melanoma (**Figure 7E and F**). Taken together, these results indicate that VPS35 regulates melanoma metastasis through MT1-MMP.

**Figure 7.**
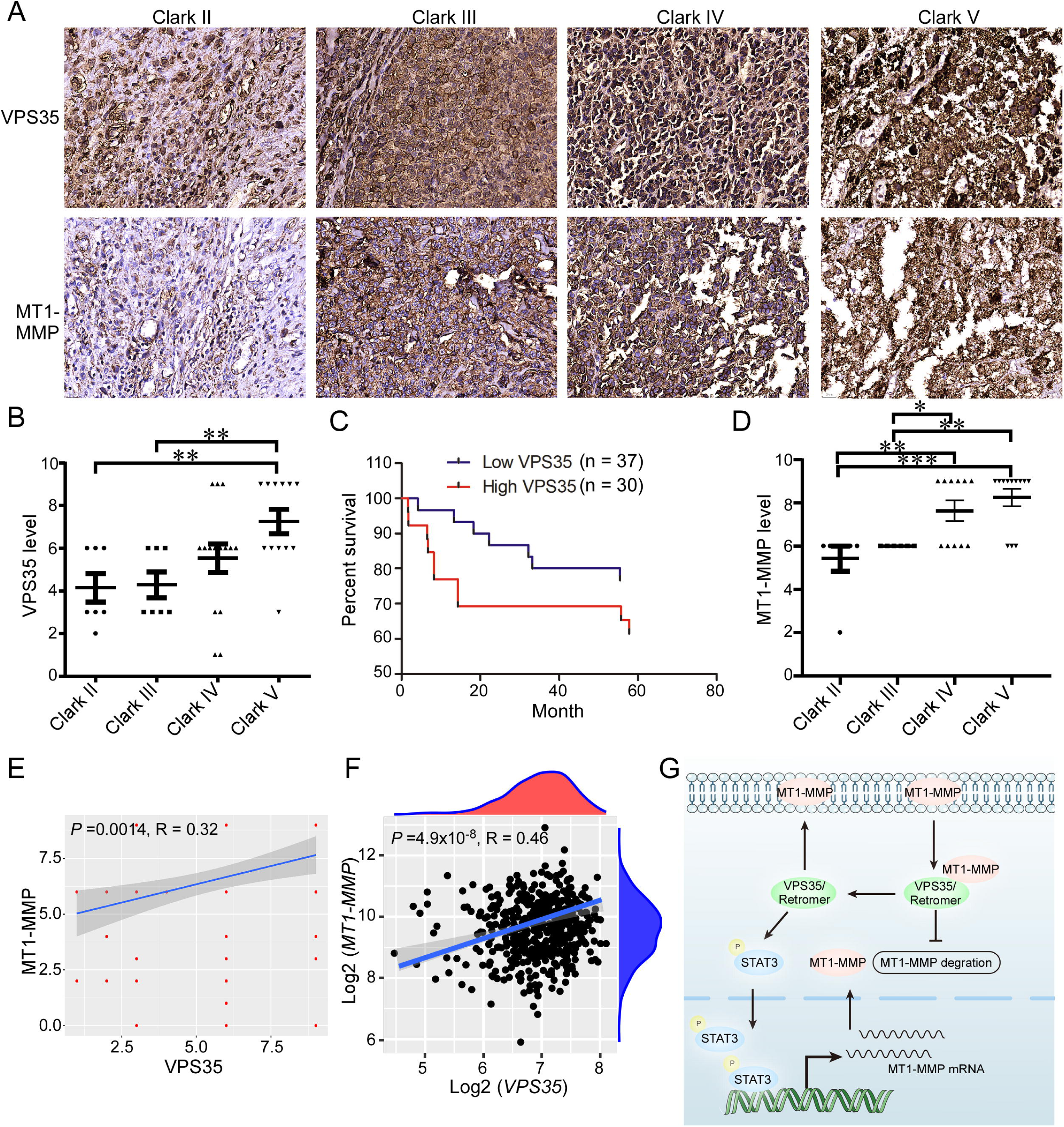
Expression of *VPS35* and *MT1-MMP* in melanoma tissues and survival analysis in melanoma patients. A. Representative images of VPS35 and MT1-MMP immunohistochemical staining across different clark grades. B. Comparative analysis of VPS35 expression levels in melanomas of different clark grades. C. Overall survival curves based on VPS35 expression in melanoma patients. D. Comparative analysis of MT1-MMP expression levels in melanomas of different clark grades. E. Correlation analysis of VPS35 and MT1-MMP expression levels in melanoma tissue. F. Correlation analysis of *VPS35* and *MT1-MMP* expression levels in the TCGA database. G. Schematic summary of VPS35/retromer in MT1-MMP regulation. MT1-MMP, as a membrane protein, is regulated in its intracellular transport and recycling by VPS35/retromer. Inhibition of VPS35/retromer restricts MT1-MMP distribution on the membrane and increases its colocalization with lysosomes, promoting MT1-MMP degradation. Furthermore, inhibition of VPS35/retromer suppresses the IL-6/STAT3 signaling pathway, reduces STAT3 nuclear translocation, and decreases *MT1-MMP* mRNA levels. In summary, VPS35/retromer regulates MT1-MMP levels through a dual mechanism, thereby influencing tumor metastasis. (\**P* < 0.05, \*\**P* < 0.01, \*\*\**P* < 0.001)

Collectively, this study reveals that VPS35/retromer has a dual effect on MT1-MMP levels by regulating MT1-MMP protein stability and promoting its transcription in melanoma. Our results highlight the critical role of VPS35/retromer in metastatic dissemination (**Figure 7G**).

## DISCUSSION

Metastasis is the primary cause of death in most cancers (Hao et al., 2021; Smalley et al., 2021). In the metastatic cascade, cancer cells spread from the primary tumor to distant organs through a series of steps: local invasion, intravasation, survival in circulation, arrest at distant sites, extravasation, and micrometastasis formation, ultimately supported by tumor angiogenesis (Valastyan and Weinberg, 2011). This process depends on cell motility, invasion, and ECM remodeling (Welch and Hurst, 2019). Malignant tumors acquire a basement membrane (BM)-breaching phenotype, allowing cancer cells to cleave adhesion molecules, breach the BM, and invade the stromal ECM. MMPs facilitate this process by degrading the BM and aiding both intra- and extravasation (Niland et al., 2021). This study highlights the VPS35/retromer’s role in melanoma metastasis, promoting tumor spread by regulating MT1-MMP protein stability and mRNA level. VPS35/retromer influences MT1-MMP protein stability through its intracellular localization and recycling, while also modulating its mRNA via the IL-6/STAT3 pathway.

*MT1-MMP* expression is associated with various malignant tumors and serves as a crucial enzyme in tumor invasion. It plays a central role in degrading the ECM during cell migration through the BM and interstitial tissues. As a key mediator of cell migration and invasion (Castro-Castro et al., 2016), *MT1-MMP* provides critical insights into cancer epigenetics and tumor migration or invasion. Targeting *MT1-MMP* could lead to new therapies aiming at reducing or halting cell proliferation and migration, potentially preventing cancers and other diseases.

The retromer complex regulates matrix invasion activity by recycling MT1-MMP. Among its components, VPS35 is the largest subunit in the retromer core, assembled with other subunits such as VPS26 and VPS29. This trimer determines cargo recognition for the recycling and degradation of target proteins (Zimprich et al., 2011). Our study discovered that VPS35 is highly expressed in melanoma patients, with elevated levels correlating with poorer prognosis. VPS35 promotes melanoma progression by regulating MT1-MMP. Tail vein metastasis experiments in nude mice demonstrated that VPS35/retromer influences pulmonary metastasis in melanoma via MT1-MMP. Silencing *VPS35* inhibits the growth of N-Ras-dependent melanoma cells (Zhou et al., 2016). However, there is limited research on the mechanism by which VPS35 dysfunction and its impact on MT1-MMP influence melanoma.

MT1-MMP is a transmembrane protein initially discovered on the cell membrane (Koziol et al., 2012). Its subcellular localization is important for its function. MT1-MMP exhibits distinct intracellular effects based on its distribution in the plasma membrane, Golgi apparatus, or cell nucleus, influencing cell movement, metabolism, and gene transcription (Koziol et al., 2012). In this study, we found that *VPS35* shRNA significantly affects the subcellular localization of MT1-MMP. In cells lacking *VPS35*, MT1-MMP’s membrane localization decreased, cytoplasmic localization increased, and it accumulated around the nucleus, leading to reduced stability and increased degradation. Previous research has shown that MT1-MMP, apart from being on the cell surface, localizes in the cytoplasm, invadopodia, Golgi apparatus, and cell nucleus, playing different roles in various intracellular processes (Laronha and Caldeira, 2020; Shimizu-Hirota et al., 2012). The structure of MT1-MMP guides its localization to extracellular and intracellular compartments, allowing for variability in its distribution(Gingras and Béliveau, 2010). We speculate that the alterations in MT1-MMP subcellular localization caused by inhibiting *VPS35* might impact melanoma in several ways: on one hand, it could reduce the degradation of the ECM around the cells, thereby inhibiting tumor cell migration and invasion; on the other hand, MT1-MMP might have different functions in various subcellular locations, such as cell migration, macrophage metabolism, invadopodium formation, spindle formation, and gene expression (Maybee et al., 2022). These functions could potentially be harnessed to inhibit melanoma progression and metastasis. Additionally, the functional loss of *VPS35* might affect MT1-MMP subcellular localization by influencing the overall function of the retromer (Collins et al., 2008; Gokool et al., 2007; Restrepo et al., 2007).

Furthermore, we report the association between VPS35 and MT1-MMP, with VPS35 regulating the lysosomal degradation pathway of MT1-MMP. VPS35 and the retromer complex play significant roles in the endolysosomal system (Williams et al., 2022). Within the endolysosomal pathway, the retromer primarily functions during the transition from early to late endosomes (Seaman, 2012). The retromer recognizes different proteins at this juncture by diverting them away from the lysosomal degradation pathway (Seaman, 2012). Disruption of the retromer leads to various cellular phenotypes, including defects in synaptic receptor recycling and alterations in lysosome-mediated degradation pathways, such as autophagy and mitophagy (Chen et al., 2019). In this study, we found that in the melanoma cell line WM793, silencing *VPS35* using shRNA promoted the association of MT1-MMP with late endosomes and lysosomes. Previous research has indicated that MT1-MMP is upregulated in human cancers and is enriched in the leading edge of invasive lesions (Lodillinsky et al., 2016; Perentes et al., 2011). In cell lines derived from breast adenocarcinomas, such as MDA-MB-231 and BT-549, a substantial portion of MT1-MMP undergoes internalization from the cell surface and accumulates in VAMP7 and RAB7-positive late endosomes/lysosomes. From there, it can be recycled to specific membrane structures involved in substrate degradation, called invadopodia(Buccione et al., 2004; Jiang et al., 2001; Monteiro et al., 2013; Uekita et al., 2001). We speculate that the impact of VPS35 on the lysosomal degradation of MT1-MMP may be related to associated lysosomal enzymes. It may be involved autophagic processes, and interact with complex components. This process might indirectly influence tumor invasion.

We found that inhibiting VPS35/retromer also affects the mRNA levels of *MT1-MMP*. VPS35/retromer not only possesses structural characteristics, but also is involved in gene regulatory mechanisms, which seem somewhat unique within retromer functionality. When *VPS35* is depleted, immune response and inflammation-related pathways are significantly enriched. Additionally, the expression of inflammatory cytokines such as IL-6 is reduced, and the nuclear translocation of STAT3 is weakened. In gastric cancer, inhibition of *VPS35* leads to reduced IL-6 expression and decreased STAT3 phosphorylation(Zhou et al., 2024). The retromer complex serves as a platform for STAT3 phosphorylation and facilitates its nuclear localization(Smalley et al., 2021). The IL-6/STAT3 signaling pathway is crucial for oncogenesis and plays a significant role in promoting cancer cell metastasis. As a transcription factor, activated STAT3 can regulate the expression of MMPs mRNA level (Hu et al., 2024; Weng et al., 2015). STAT3 activity can regulate MT1-MMP expression (Weng et al., 2015). In our results, depleting *VPS35* and overexpressing STAT3 significantly restored *MT1-MMP* levels. Together, these studies indicate the critical role of VPS35 in regulating *MT1-MMP* mRNA levels through the IL6/STAT3 pathway.

VPS35/Retromer regulates tumor metastasis via MT1-MMP. In a melanoma lung metastasis model, VPS35-deficient melanoma cells formed fewer lung metastases, while MT1-MMP expression accelerated metastasis. Similarly, melanoma tissue microarrays revealed high VPS35 and MT1-MMP levels correlated with advanced staging and poor prognosis. These findings underscore the pivotal role of VPS35/retromer in promoting metastasis through MT1-MMP.

Our findings reveal that VPS35/retromer mediates melanoma metastasis by enhancing MT1-MMP protein stability through recycling and increasing its mRNA levels via the STAT3 pathway. Our study confirms the molecular mechanisms by which VPS35/retromer regulates MT1-MMP levels. Targeting the VPS35/retromer-MT1-MMP regulatory pathway presents a novel therapeutic strategy for cancer metastasis treatment, thus offering new avenues for combating tumor metastasis.

## MATERIALS AND METHODS

### Antibodies

Rabbit antibodies against VPS35 (R27432, ZEN-BIOSCIENCE, 1:1000 for immunoblots (IB)), MT1-MMP (R22533, ZEN-BIOSCIENCE, 1:1000 for IB, 1:100 for PLA), H3 (ET1701-64, HUABIO, 1:1000 for IB), STAT3 (10253-2-AP, proteintech, immunoprecipitation (IP): 4 μg for 3 mg of total protein lysate). Mouse antibodies against VPS35 (B-5, sc-374372, Santa Cruz Biotechnology, IP: 2 μg for 500 μg of total protein lysate, 1:200 for PLA), STAT3 (sc-8019, Santa Cruz Biotechnology, 1:1000 for IB), p-STAT3 (sc-8059, Santa Cruz Biotechnology, 1:1000 for IB), GAPDH (sc-32233, Santa Cruz Biotechnology, 1:1000 for IB), SNX27 (ab77799, Abcam, 1:1000 for IB).

### Cell culture

WM793, 1205Lu, SW480 and SW620. The cell culture media used were RPMI 1640 (for all cells). All cell culture media were supplemented with 10% FBS and Levofloxacin. For all the cells used, mycoplasma contamination was checked.

### Plasmids and cell transduction

Lentiviral shRNA plasmids targeting *VPS35* and *STAT3* were cloned into the pLKO.1-TRC vector (select with puromycin).

Target sequence as flow:

*VPS35* shRNA1 target sequence: GCAGATCTCTACGAACTTGTA;
*VPS35* shRNA2 target sequence: CAGTGAAGAAACAGAGCAGAT;
*STAT3* shRNA target sequence: GCACAATCTACGAAGAATCAA.

The *SNX27* lentiviral shRNAs plasmid was supplied by Chonglin Yang Lab (Kunming, China). The human *MT1-MMP* CDS was cloned into pLNCX2-GFP vector (select with G418). The human *STAT3* CDS was cloned into the pLenti-cmv-blast vector (select with blasticidin). For lentivirus or retroviral production, HEK-293T cells were transfected with the respective vector, packaging plasmid psPAX2 (gag/pol for retroviral), and VSVG using 1 mg/mL polyethyleneimine, following the manufacturer’s instructions. Viruses were harvested 48–72 hours post-transfection, and target cells were transduced with the lentivirus. After 48 hours, resistant cells were selected using puromycin, G418 or blasticidin before additional experiments were performed.

### RT-qPCR assay

Total RNA was extracted using the TRIZOL method, and cDNA was synthesized using the First-strand cDNA Synthesis Mix Kit (F0202, LABLEAD). qPCR was performed in an iCycler thermocycler (Bio-Rad Laboratories) using the 2× Relab Green PCR Fast Mixture (Universal) Kit (R0202-02, LABLEAD). mRNA levels were quantified using iCycler software (Bio-Rad Laboratories) and normalized to GAPDH. The primers used for RT-qPCR were as follows:

*MT1-MMP* q-PCR NS: 5’-TGCCCAATGGAAAGACCTAC-3’
*MT1-MMP* q-PCR CAS: 5’-GTACTCGCTATCCACTGCC-3’

### Western blot assay

Cells were scraped and lysed in RIPA buffer on ice for 10 minutes. Then, 30 μg of total protein was loaded onto SDS-PAGE gels. The gels were initially run at 60 V for 30 minutes, followed by 120 V until the dye front exited the gel. Proteins were transferred to PVDF membranes, which were then blocked with 5% non-fat dry milk in Tris-buffered saline containing 0.05% Tween-20 (TBS-T) for 30 minutes at room temperature. After blocking, the membranes were incubated with primary antibodies for 2-4 hours at 4°C, followed by a 60-minute incubation with secondary antibodies at room temperature. Detection was performed using an ECL-plus western blotting detection system (Tanon-5200Multi).

### Membrane protein biotinylation assay

Membrane protein biotinylation assay WM793 cells grown to 90% confluence were washed twice with icecold PBS, and freshly prepared 1 mM EZ-link sulfo-NHS-SS-biotin (Thermo Fisher Scientific) in ice-cold PBS was added. After incubation on ice for 30 min, cells were washed twice with ice-cold PBS. The unreacted biotin was quenched for 10 min by incubation with ice-cold PBS containing 50 mM glycine. Cells were then washed three times with ice-cold PBS and lysed in 0.5 mL IP buffer (PBS, pH 7.4, 5 mM EDTA, 5 mM EGTA, 10 mM sodium pyrophosphate, 50 mM NaF, 1 mM NaVO_3_, and 1% Triton X-100). Cell lysate was briefly sonicated (5 s) and centrifuged at 14,000 rpm on a 4°C bench top centrifuge. The supernatant was incubated with Streptavidin Magnetic Beads at 4°C for overnight. The beads were extensively washed with IP buffer, eluded with SDS-PAGE loading buffer, and subjected to western blotting.

### Immunofluorescence staining

Melanoma cells (3 × 10^4^) in RPMI 1640 growth medium were plated onto gelatin-coated glass coverslips for 12 h and fixed with 4% paraformaldehyde. The cells were then permeabilized in antibody diluting buffer (2% BSA and 0.1% Triton X-100 in PBS) and incubated with the indicated antibodies, for 2 h. The cells were incubated with the secondary antibodies conjugated with Alexa Fluor 488 or Alexa Fluor 594 dye for 1 h at room temperature before confocal microscopy. An extensive wash was performed between each step. The coverslips were then mounted onto slides and imaged using laser scanning confocal microscope.

### Proximity ligation assay

Proximity ligation assay (PLA) was performed using the Duolink® In Situ kit (Sigma-Aldrich) to investigate the interaction between VPS35 and MT1-MMP in fixed cell samples. Cells were cultured on glass slides, fixed with 4% paraformaldehyde (PFA) for 15 minutes at room temperature, and permeabilized with 0.1% Triton X-100. Samples were blocked using the Duolink blocking solution to prevent nonspecific binding, followed by incubation with primary antibodies against VPS35 (mouse) and MT1-MMP (rabbit) overnight at 4°C. For the negative control, VPS35 antibody was replaced with an equivalent amount of mouse IgG to assess nonspecific interactions. After washing with filtered PBS, the samples were incubated with species-specific Duolink PLA probes, and the samples were incubated in a humidity chamber at 37°C for 1 hour, and were incubated with ligation–ligase solution (30 min, 37 °C) in a pre-heated humidity chamber. After washing, coverslips were incubated with the amplification reaction mixture (100 min, 37 °C) in a pre-heated humidity chamber, washed, and counterstained with DAPI. The images were observed by a confocal microscope (Zeiss, Germany) and analyzed by the ZES software. Each green dot represents the detection of protein–protein interactions.

### Co-immunoprecipitation (Co-IP) assay

Cells were harvested in IP lysis buffer (50 mM Tris, pH 8.0, 150 mM NaCl, 1% Triton X-100, 0.5% Sodium deoxycholate) with 1 mM phosphorylase inhibitor, incubated on ice for 10 minutes, ultrasonic cell disruption, and centrifuged at 12,000 × g for 10 minutes. The antibody (MT1-MMP, VPS35, STAT3, or IgG control) was incubated with total protein at 4°C for 3 hours, followed by the addition of Protein A/G agarose beads and further incubation overnight at 4°C. Complexes were washed, eluted, boiled for 10 minutes, and analyzed by western blot.

### *In vivo* metastasis assays

All animal procedures followed protocols approved by the Laboratory Animal Center of Yunnan University. Nude mice were used for experimental lung metastasis experiments. 1205Lu human melanoma cells expressing the luciferase reporter were trypsinized and washed with PBS. Subsequently, 1×10^6^ cells in 0.2 mL PBS were injected into the lateral tail vein. Luciferase-based noninvasive bioluminescent imaging and analysis were performed as described previously with an imaging system (Tanon ABL X5 system) by injecting 100 µL d-luciferin (15 mg/mL) (Hao et al., 2024). The mice were imaged on day 0 and then on a weekly basis thereafter. At the end of the metastasis experiment, lungs were harvested from euthanized mice, fixed in paraformaldehyde, and embedded in paraffin. Paraffin sections were stained with hematoxylin and eosin.

### Statistical analysis

Statistical analysis was performed using an unpaired Student’s t test in GraphPad Prism. *P*[<[0.05 was considered to indicate statistical significance.

## Compliance and ethics

The authors declare no competing financial interests.

## Acknowledgement

This work was supported by National Natural Science Foundation of China (82273460, 32260167), the Yunnan Fundamental Research Projects (202401AS070133), a grant (KLTIPT-2023-02) from Key Laboratory of Tumor Immunological Prevention and Treatment in Yunnan Province, Yan’an Hospital Affliated to Kunming Medical University, and grants (KC-23233927, ZC-24249243) from Yunnan University. We thank Dr. Jing Li and Shengyu Yang for critical reading of the manuscript.

