## Supplemental Figure for "VPS35/Retromer-dependent MT1-MMP regulation confers melanoma metastasis"

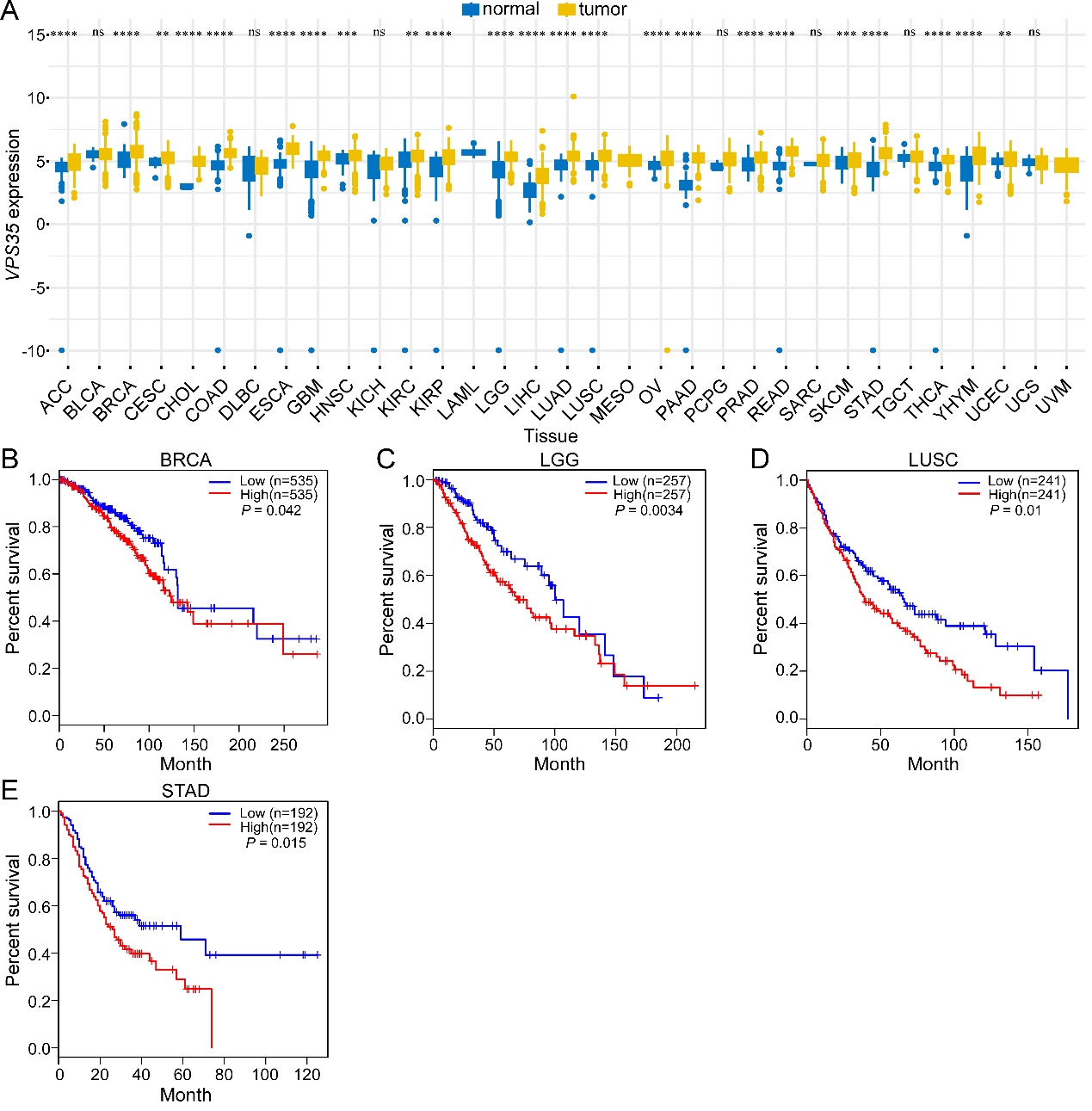

**Figure S1. Expression of *VPS35* in the TCGA and GTEx databases.** A**.** Expression of *VPS35* across different tumor types in the TCGA and GTEx databases. **B-E** Survival analysis of *VPS35* in breast invasive carcinoma (B), brain lower grade glioma (C), lung squamous cell carcinoma (D) and stomach adenocarcinoma (E).

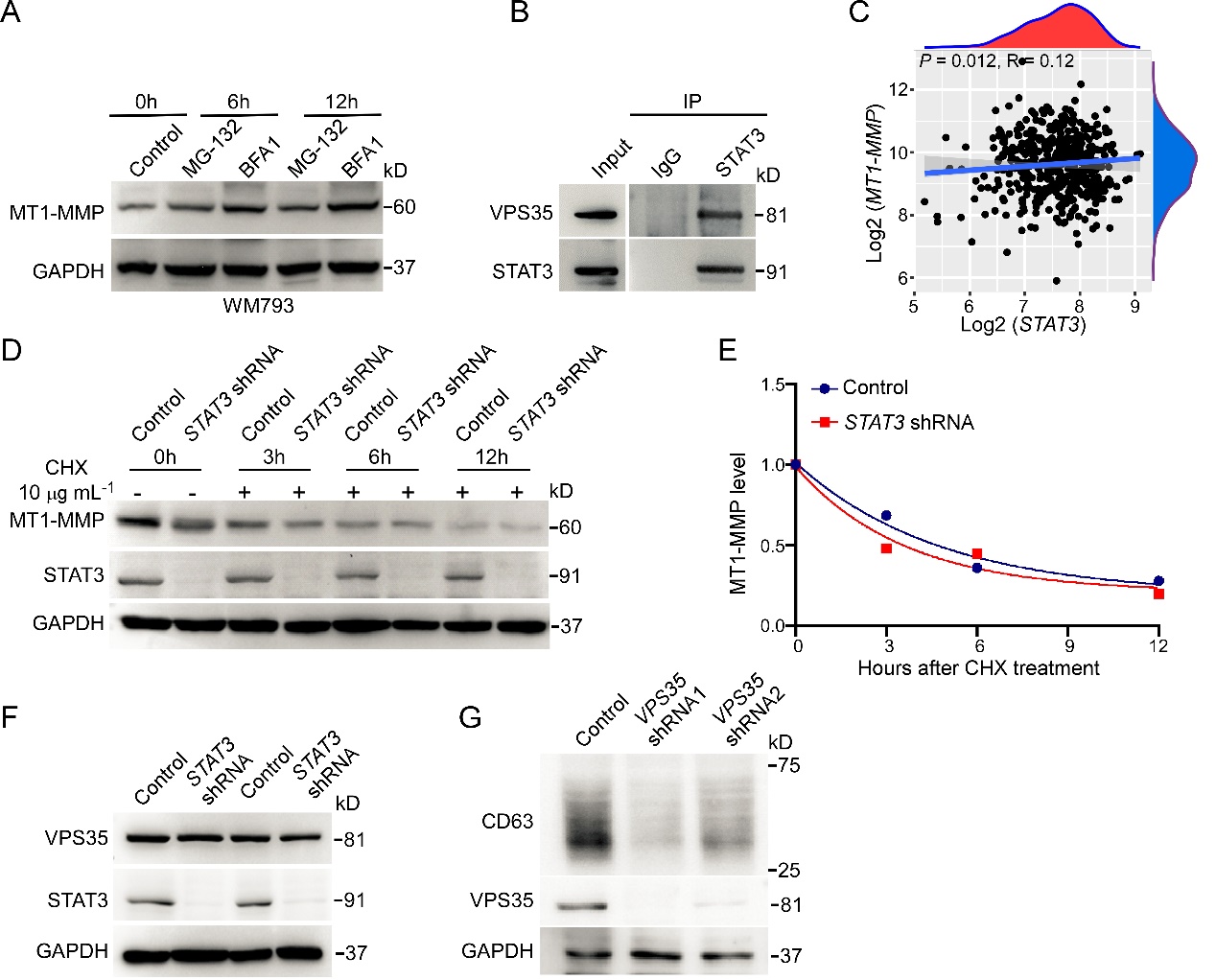

**Figure S2. STAT3 interacts with VPS35 and correlates significantly with *MT1-MMP* expression in melanoma.** A**.** Western blotting analysis of MT1-MMP levels following 5 μM MG-132 and 100 nM BFA1 treatment in WM793 wild-type cells. B. Co-IP analysis showing that VPS35 interacts with STAT3. C. Correlation analysis of *STAT3* and *MT1-MMP* expression levels in the TCGA database. D. Western blotting analysis of MT1-MMP stability following 10 μg/mL CHX treatment in WM793 control and *STAT3* shRNA cells. E. Quantitative analysis of MT1-MMP stability in WM793 control and *STAT3* shRNA cells. F. Western blotting analysis of VPS35 level in *STAT3* shRNA cells. G. Western blotting analysis of CD63 levels in WM793 control and *VPS35* shRNA.
